# Context dependent isoform specific PI3K inhibition confers drug resistance in Hepatocellular carcinoma cells

**DOI:** 10.1101/2021.03.17.435839

**Authors:** Kubra Narci, Deniz Cansen Kahraman, Altay Koyas, Tulin Ersahin, Nurcan Tuncbag, Rengul Cetin Atalay

**Affiliations:** Cancer System Biology Laboratory, CanSyL, Graduate School of Informatics, Middle East Technical University, 06800, Ankara, Turkey; Section of Pulmonary and Critical Care Medicine, the University of Chicago, 60637, Chicago, IL, USA

**Keywords:** Liver Cancer, PI3K/Akt/mTOR Pathway, Network Analysis, Synergy, Resistance

## Abstract

**Background:** Targeted therapies for Primary liver cancer (HCC) is limited to the multi-kinase inhibitors, and not fully effective due to the resistance to these agents because of the heterogeneous molecular nature of HCC developed during chronic liver disease stages and cirrhosis. Although combinatorial therapy can increase the efficiency of targeted therapies through synergistic activities, isoform specific effects of the inhibitors are usually ignored. This study concentrated on PI3K/Akt/mTOR pathway and the differential combinatory bioactivities of isoform specific PI3K-*α* inhibitor (PIK-75) or PI3K-*β* inhibitor (TGX-221) with Sorafenib dependent on PTEN context.

**Methods:** The bioactivities of inhibitors on PTEN adequate Huh7 and deficient Mahlavu cells were investigated with real time cell growth, cell cycle and cell migration assays. Differentially expressed genes from RNA-Seq were identified by edgeR tool. Systems level network analysis of treatment specific pathways were performed with Prize Collecting Steiner Tree (PCST) on human interactome and enriched networks were visualized with Cytoscape platform.

**Results:** Our data from combinatory treatment of Sorafenib and PIK-75 and TGX-221 showed opposite effects; while PIK-75 displays synergistic effects on Huh7 cells leading to apoptotic cell death, Sorafenib with TGX-221 display antagonistic effects and significantly promotes cell growth in PTEN deficient Mahlavu cells. Transcriptomic states of combinatory treatments with PIK-75 and TGX-221 inhibitors were identified in PTEN deficient and adequate cells. Molecular interactions and cell signaling pathways were reconstructed and analyzed in-depth to understand mechanism of differential synergistic or antagonistic effects of PI3K-*α* (PIK-75) and PI3K-*β* (TGX-221) inhibitors with Sorafenib.

**Conclusions:** Simultaneously constructed and analyzed differentially expressed cellular networks presented in this study, revealed distinct consequences of isoform specific PI3K inhibition in PTEN adequate and deficient liver cancer cells. We demonstrated the importance of context dependent and isoform specific PI3K/Akt/mTOR signaling inhibition in drug resistance during combination therapies. (https://github.com/cansyl/Isoform-spesific-PI3K-inhibitor-analysis).

## Background

According to WHO-Global cancer observatory (GCO) that one-fifth of men and one-sixth of women will diagnosed to cancer throughout their lives and one-eighth of men and one-eleventh of women will die of it worldwide before the age of 75 years. Hepatocellular cancer (HCC) which constitutes the 75% of Primary liver cancers is the 5th most common and the 3rd most lethal cancer in the world [1, 2]. While the the death rates from other cancers are decreasing due to advances in diagnosis and therapeutics, the incidence and the mortality of HCC follow an increasing trend due to high rate of obesity associated liver diseases [3, 4].

Development of HCC is multi-factorial and complex biological process, where the chronic liver disease is initiated due to hepatic injury, followed by the continuous inflammation and cell death, which in turn leads to the regeneration of hepatocytes and the increased rate of mutations along with genomic instability [5]. The increased number of proliferating cells evokes the activation of several cell signaling pathways involved in liver regeneration, such as growth factor signaling, cell differentiation, angiogenesis and cell survival. Stimulation of these pathways is mostly associated with tyrosine kinases which are usually the members of PI3K/Akt/mTOR cell signaling [6]. Studies show that the heterogeneous nature of HCC is mostly caused by the variations of mutations and alterations in expression levels of these key proteins [7].

Currently there is no effective therapy for patients suffering from HCC, the survival rates is only 7% for five years [1]. There are two FDA approved small molecule drug treatments for HCC; Sorafenib (Nexavar, BAY43-9006) and Regorafenib (Bayer, BAY73-4506), are receptor tyrosine kinase inhibitors targeting Raf, VEGFR and PDGFR kinases. They inhibit tumor cell proliferation and angiogenesis while promoting apoptosis. However, in most of the cases they are not capable of eliminating the cancer cells mostly because of the heterogeneous nature of HCC [8, 9]. Moreover, the signaling pathways involved in proliferation, growth, angiogenesis and metastasis are redundant, compensating each other though some key molecular regulations. Which makes them with superfluous functions due to the potential cross-talks between them, which could be another reason for the ineffectiveness of these two multi-kinase inhibitors [6].

The constitutive activation of PI3K/Akt/mTOR signaling pathway is frequently observed in liver cancer due to inactivating mutations or loss of heterozygosity in a tumor suppressor protein Phosphatase and tensin homolog (PTEN). PTEN disfunction is observed in nearly 50% of the HCC cases and correlated with poor prognosis, drug resistance and low patient survival [10, 11, 12]. PTEN prevents the Akt activation by dephosphorylating PIP3, or mutations activating PIK3CA gene, or damage in the negative-feedback loop from mTOR signaling pathway in various epithelial cancers including HCC [13, 14, 15, 16].

The influence of isoform diversity on responses to drugs with respect to large number of GPCR receptors is demonstrated at systems level recently [17]. Furthermore there are resent studies on the association of isoform specific differential involvement of AKT in the pathophysiology and therapeutic responses of cancer cells [18, 19, 20]. Here in this study we focused on the response of HCC cells isoform specific PI3K inhibitors. PI3Ks grouped into three classes based on their structures [21, 22] but two of Class I members of PI3Ks have heterodimeric class IA p110-*α* (p110) and class IB p110-*β* (p85) regulatory subunits are well studied enzymes in cancer. PIK3CA gene encoded PI3K isoform p110-*α*, is activated through receptor tyrosine kinases (RTKs) and Ras oncogene. In cancer, signaling though PI3K predominantly depends on alpha isoform regulating cellular growth, metabolism and angiogenesis. The other PI3K isoform encoded by PIK3CB, p110-*β* is regulated mostly by G protein-coupled receptors (GPCRs) and has critical functions in inflammatory cells [23, 24].

In this study we demonstrated context (PTEN function) dependent isoform specific PI3K inhibition confers drug resistance by their antagonistic and synergistic effects with Sorafenib on HCC cells at network level in and studies focusing on the discovery of agents against HCC aim to identify target proteins that escape from regulatory signaling mechanisms of the cell.

## Results

### Molecular and cellular characterization Huh7 and Mahlavu cells in the presence of small molecule isoform specific PI3K inhibitors

Well-differentiated Huh7 cell line with adequate PTEN and poorly-differentiated PTEN deficient Mahlavu cells were selected to exploit throughout this study. The expression levels and the phosphorylation status of key proteins in PI3K/Akt/mTOR and RAF/MEK/ERK signaling pathways were reported by our group, and in correlation with their PTEN status, Mahlavu cells display hyper-activated cell survival proteins [25]. Initially Sorafenib, LY294002, PI3K inhibitor p110*α* subunit specific (PIK-75) and PI3K inhibitor p110*β* subunit specific (TGX-221) were analyzed for their cytotoxic bioactivity and their effect on cell cycle progression with Huh7 and Mahlavu (Fig. 1A). G1, S and G2/M cell cycle phases were analyzed separately to calculate viable cell distributions among them (Fig. 1B). Sub-G1 percentage demonstrating apoptotic cells were also calculated. Cell cycle distribution remain stable for both cell lines and all inhibitor treatments. In both cell lines, Sorafenib and PIK-75 treatments showed stimulation of apoptosis through increase in sub-G1 population. In Huh7, Sorafenib seems to be more active while PIK-75 functioned more in Mahlavu cells which was more aggressive than Huh7 cell line by PTEN-loss based hyper-active Akt stimulation.

**Figure 1.**
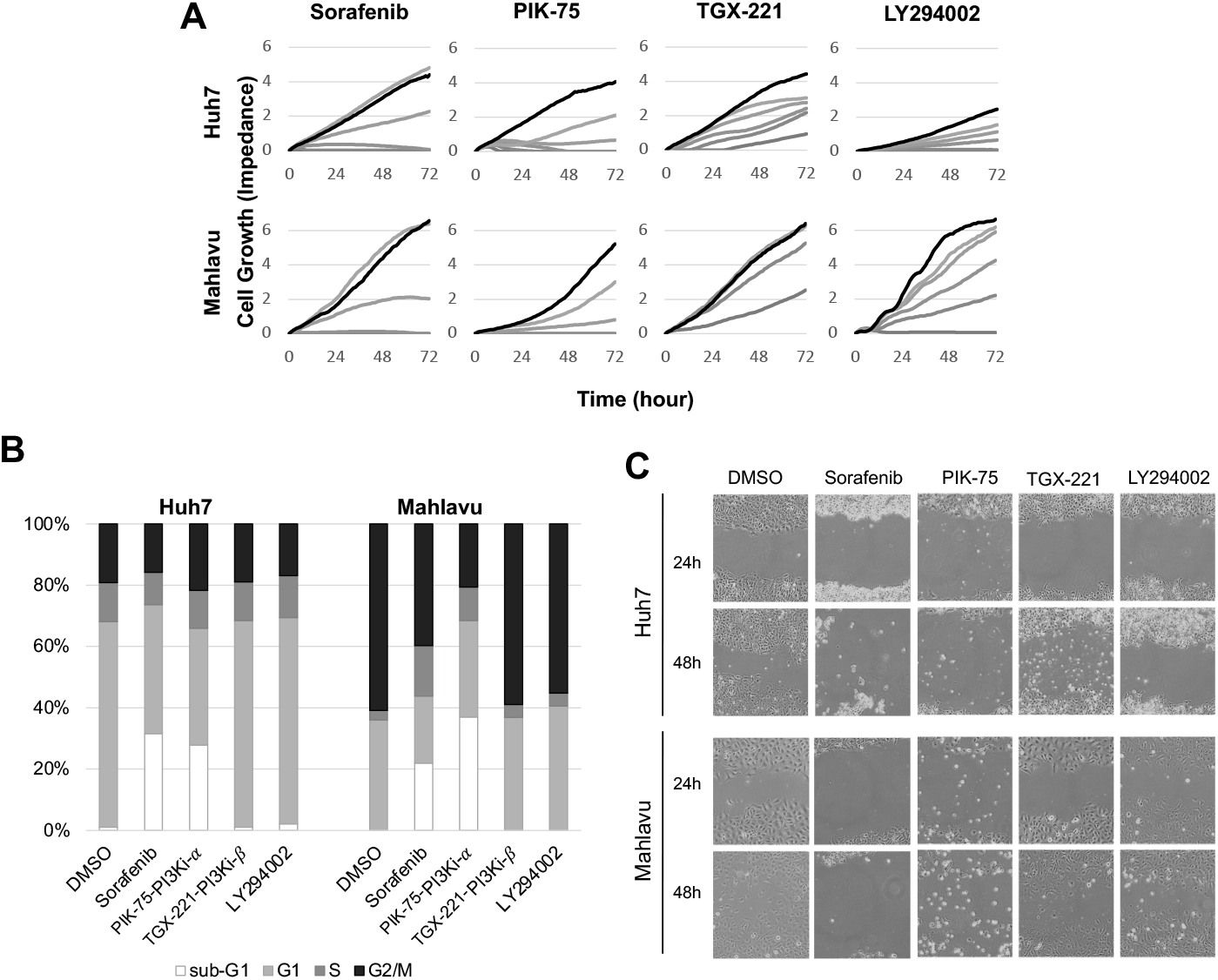
Characterization of HCC cells in the presence of small molecules inhibitors. Real time cell growth analysis of Huh7 and Mahlavu cells with increasing concentrations (40*μ*M, 20*μ*M, 10*μ*M, 5*μ*M, 2.5*μ*M) of Sorafenib, PI3K inhibitor LY294002, PI3Ki-*β* inhibitor (TGX-22) and PI3Ki-*α* (1*μ*M, 0.5*μ*M, 0.25*μ*M, 0.125*μ*M, 0.0625*μ*M) PI3Ki-*α* (PIK-75) along with DMSO vehicle control (Control is black and increasing drug concentrations is given in grey level, highest concentration is being the darkest) **(A)**. Cell cycle analysis with flow cytometry. Sub-G1 population represents apoptotic cells **(B)**. Wound healing assay for 24 and 48 hours for cell migration. **(C)**. 10*μ*M of Sorafenib, LY294002 and PI3Ki-*β* (TGX-221) and 0.1*μ*M of PI3Ki-*α* (PIK-75) were used for cell cycle and migration assays.

### Migration analysis of the inhibitors

In order to analyze the effects of selected inhibitors on cell migration, wound-healing assay was performed. The percentages of wound closures after 48 hours of initial scratch were calculated for Huh7 and Mahlavu. We observed that Sorafenib and PIK-75 reduced migration significantly (p < 0.001) in both Huh7 and Mahlavu (Fig. 1C).

### Synergistic cytotoxicity analysis

Since none of the treatments alone was fully effective to inhibit growth and stimulate apoptosis, we addressed the value of co-treatments of Sorafenib with PIK-75 and TGX-221 through real-time cell growth analysis (Fig. 1C). A synergistic effect of Sorafenib and PIK-75 treatments was observed on growth of both cell lines. TGX-221 combinatory treatment with Sorafenib also resulted in synergistic growth inhibition on Huh7 cell line. On the other hand, TGX-221 displayed a growth inhibition of Mahlavu, TGX-221 co-treatment with Sorafenib resulted in an antagonistic effect and stimulated cellular growth. Furthermore, Sorafenib and PIK-75 treatment had more drastic effect on Mahlavu compared to Huh7. Therefore, these findings indicate that in PTEN deficient Mahlavu cells, constitutive activation of PI3K/Akt signaling mainly depend on p110-*α* (Fig. 2).

**Figure 2.**
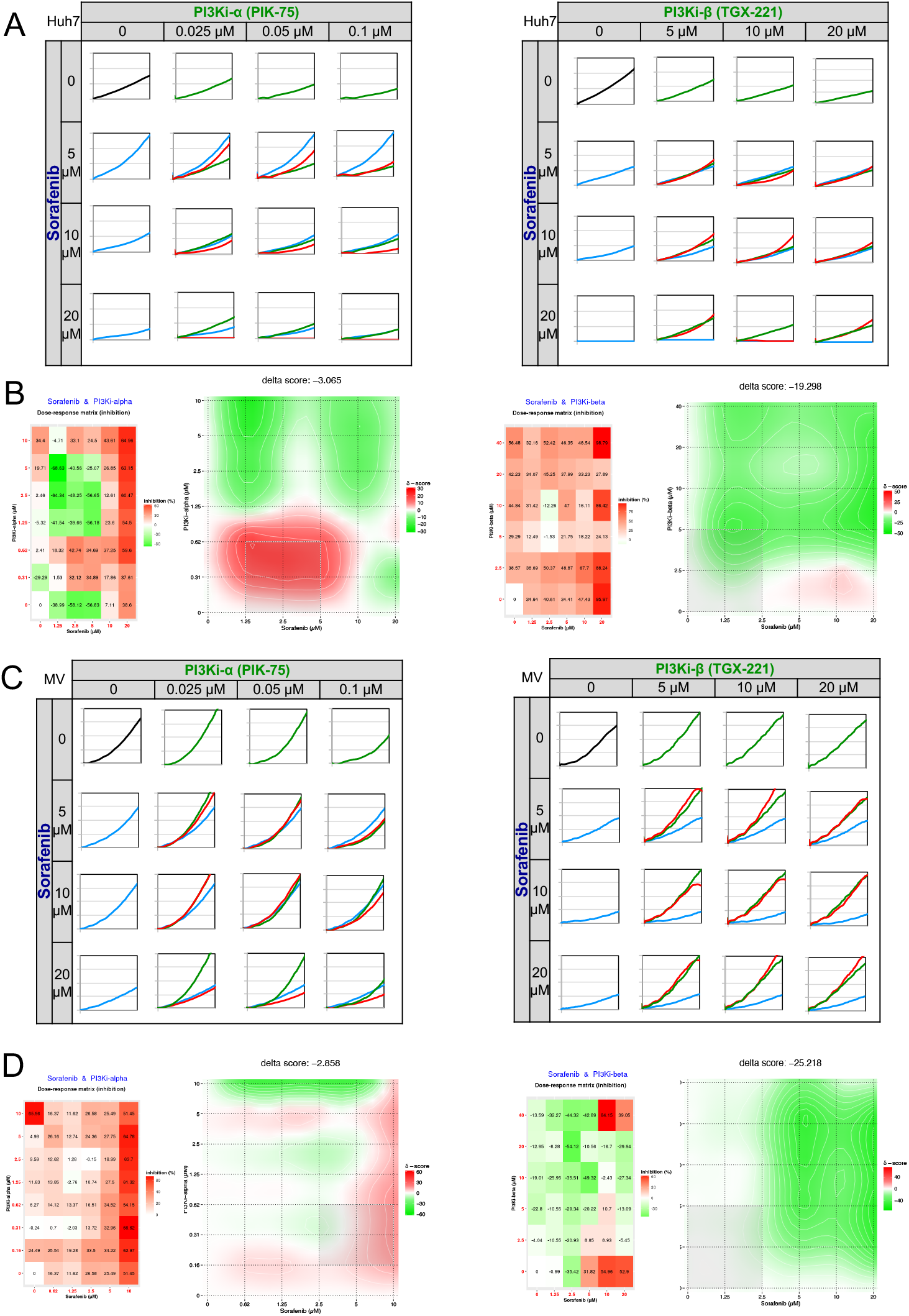
Real-time cell growth analysis. Human liver cancer cells Huh7 **(A,B)** and Mahlavu (MV) **(C,D)** were treated with the Sorafenib, PI3Ki-*α* and PI3Ki-*β* alone or in combination with increasing concentrations as indicated. Cell index measurements were obtained by RT-CES software. DMSO was used as negative control **A B**. 72 hours of the percent growth inhibition values were used to calculate drug interactions with The SynergyFinder web application. Positive delta score reflects synergistic and negative score reflects antagonistic drug interactions. Experiments were performed in triplicate.

### Network level analysis of isoform specific combinatory effects of PI3Ks

In order to describe the molecular events in the differential response of PTEN adequate (Huh7) and deficient (Mahlavu) cells displaying differential PI3K/Akt/mTOR pathway activities toward PI3K-*α* inhibitor (PIK-75) and PI3K-*β* inhibitor (TGX-221) alone or in combination with Sorafenib, we performed RNA sequencing experiments. Further, network based data analysis using systems biology approaches which is represented in Fig. 3A are applied.

**Figure 3.**
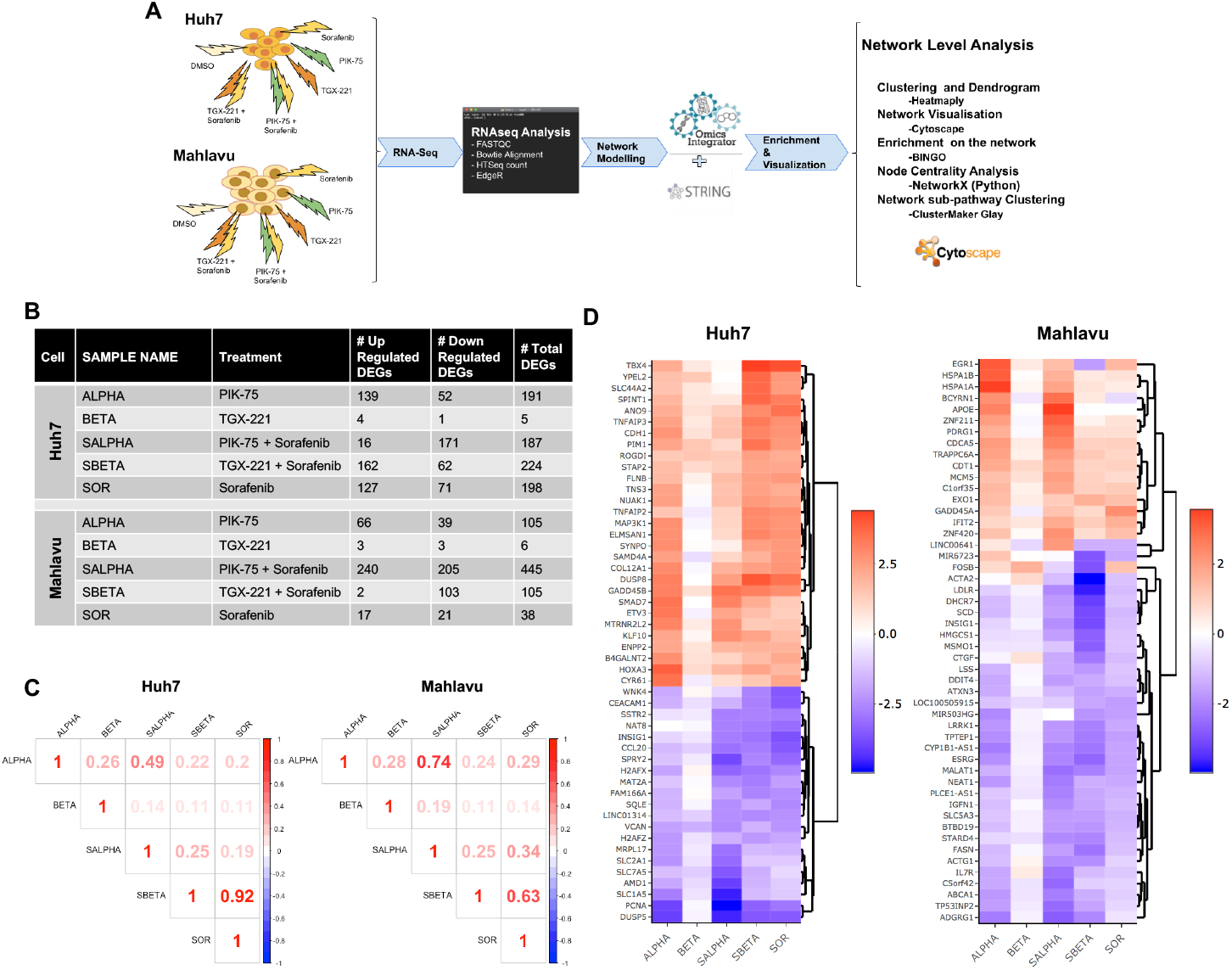
Systems Biology flowchart for RNA-seq Data analysis. Systems level methodology flowchart for differential PIK3K/Akt/mTOR pathway activities in Huh7 and Mahlavu calls treated with Sorafenib, PI3K-*α* inhibitor PIK-75 and PI3K-*β* inhibitor TGX-221 alone or in combination **(A)**. Differentially Expressed Genes (DEG) Table summarize the abbreviations of samples as the treatments to HCC cells and differentially expressed gene (DEG) numbers. DEG filtration for A and B as follows; Huh7 cells:logFC ≥ 2.0, ≤ −2.0 and p ≤ 0.01, Mahlavu cells: logFC ≥ 1.5, ≤ −1.5 and p ≤ 0.01 **(B)**. Pearson correlations for gene expressions in Huh7 and Mahlavu, no filtration**(C)**. Dendrogram analysis on logFC for top 50 DEGs, red and blue color represented for up- and downregulated genes **(D)**.

Initially we identified differentially expressed genes (DEGs) by using empirical analysis of digital gene expression data in R (edgeR tool) [26]. While PIK-75 (ALPHA) treated Huh7 cells had more upregulated genes, combination with Sorafenib reversed the number of genes in favor of gene downregulation. The response of Mahlavu to the same combination treatment (SALPHA) is significantly different with highly increased number of up and downregulated genes when compared to Sorafenib or PIK-75 alone. In both cell lines, TGX-221 treatment had minor action yet in PTEN deficient Mahlavu, single treatment of Sorafenib had a neutral effect. Interestingly, TGX-221 and Sorafenib combination resulted in higher number of downregulated genes in Mahlavu comparing the number of upregulated genes in Huh7 (Fig. 3B).

Using Pearson correlation analysis, we demonstrated the similarities in the overall gene expressions using the corresponding logFC values. A significant correlation was observed between Sorafenib alone treatment and its combinatory treatment with TGX-221 (0.92) in Huh7 and (0.63) in Mahlavu indicating ineffectiveness of single TGX-221 treatments in both cells. The similarity between PIK-75 and its combinatory treatment with Sorafenib in Mahlavu (0.74) is also high (Fig. 3C) may be an evidence of underfill action of Sorafenib alone treatment in PTEN deficient cells.

50 most commonly differentially regulated genes ranked by sum of absolute logFC values were represented through a dendrogram in Fig. 3D. In Huh7, up- and down-regulated genes were well clustered. DUSP5, PCNA, VCAN, GADD45B and DUSP8 genes in Huh7 were the shared mostly. In Mahlavu, DEGs were not well separated like Huh7, some of the genes like EGR1, LINC00641, MIR6723, FOSB and ACTA2 were found to vary in different treatments. ESRG, CYP1B1-AS1, LDLR and TPTEP1 genes were the most common DEGs in Mahlavu.

### Gene enrichment analysis of differential expression patterns in HCC cells

Considering the high correlation between specific treatments, in order to investigate the expression patterns between different inhibitory treatments in HCC cell lines, Huh7 and Mahlavu genes were clustered separately using their corresponding logFC values.

Heatmap analysis of DEGs revealed expression pattern of HCC cells and the gene enrichment analysis to these patterns (gene clusters) exploit the functional consequences (Fig. 4). In Huh7 cell line, expressions of single Sorafenib and its combined treatment with TGX-221 were highly correlated, which was also visualized in heatmap analysis. For all treatments in HCC, a positive regulation of extracellular matrix organization and developmental processes were observed while regulation of cell proliferation and actin filament bundle assembly ontologies were more active in single PIK-75 treatment. PIK-75 and Sorafenib combined treatment resulted in downregulation of genes enriched in negative regulation of biosynthetic processes and cell fate commitment ontologies. Likewise, cholesterol metabolic process was downregulated for TGX-221 and its Sorafenib combinatory treatment. We also identified a group of genes involved in apoptosis stimulation process, the genes are 2 histone family proteins, 1 long intergenic non-translating RNA, uncharacterized proteins FAM184B and NCBP2AS2, NBP and NAG5 and ATP2A1 downregulated in the treatment of PIK-75 alone while they were upregulated all the other Huh7 treatments.

**Figure 4.**
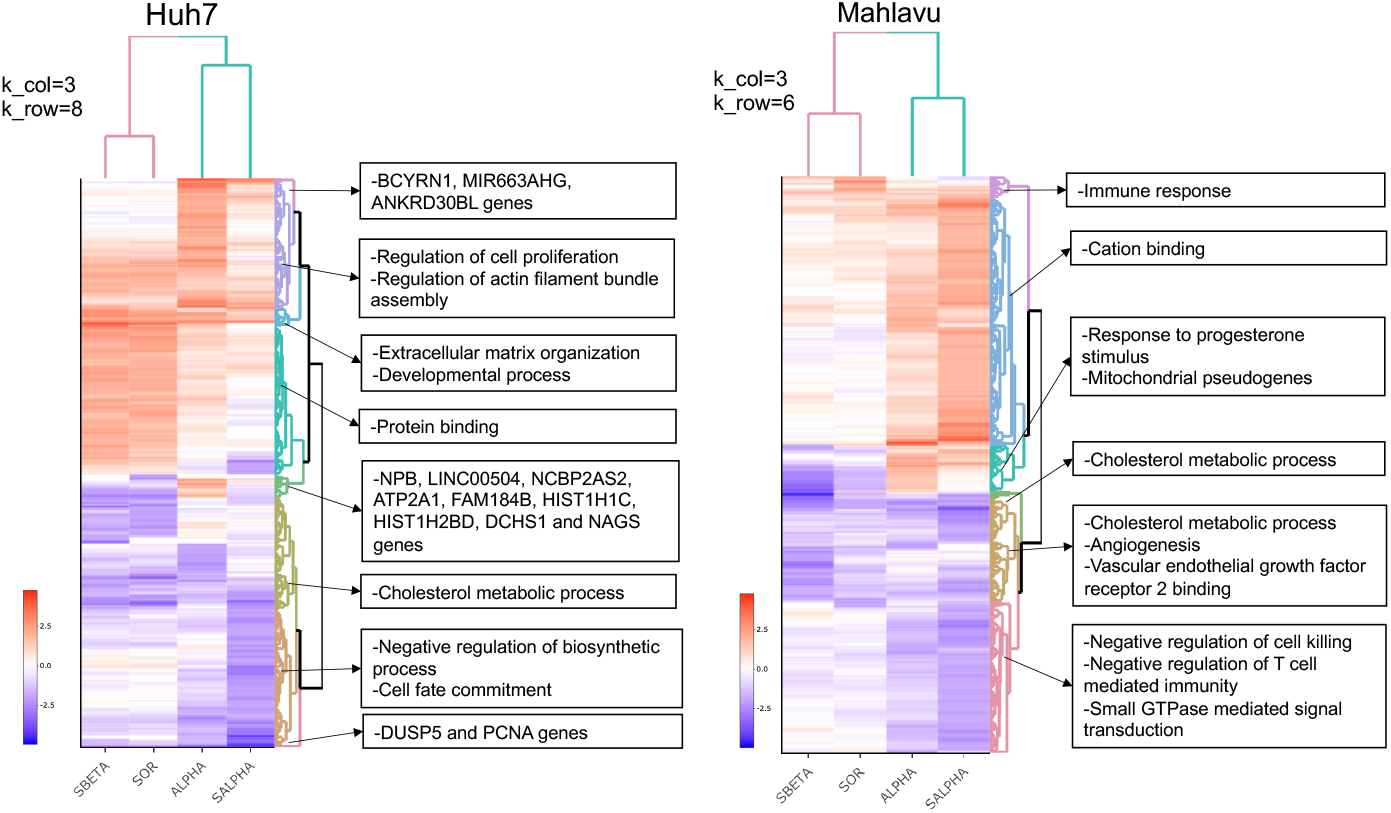
Gene Expression Patterns. Heatmaps of gene expressions illustrated as dendrograms separately for Huh7 and Mahlavu cells lines. We removed single PI3K-*β* inhibitor treatments for both cell lines considering its ineffectiveness. Sample sets for Huh7 and Mahlavu were separately joined, and united sets included 11033 and 11615 genes in total before filtration. Gene enrichment analysis was performed using BiNGO (FDR ≤ 0.05) and significant gene ontologies were selected according to the context. Hence, dendrogram analysis were performed on 581 genes for Huh7 and 583 genes for Mahlavu cells. For more detailed analysis and to view interactive dendrogram please see CanSyl github repository. Clusters were generated by heatmaply and colored; 8 for Huh7 and 6 for Mahlavu. Clusters not showing any significant enrichment were excluded. Up- and downregulated gene expression levels are colored as red and blue respectively, the intensity of the color indicates how strong the logFC value is. ALPHA; PIK-75, SALPHA; PIK-75 and Sorafenib, SBETA; TGX-221 and Sorafenib, SOR; Sorafenib treatments.

Immune response was downregulated more significantly in Mahlavu cells in combined with Sorafenib treatment. Cation binding was enhanced for PIK-75 and its Sorafenib combination. Cholesterol metabolic processes, angiogenesis and vascular endothelial growth factor receptor 2 binding were downregulated for all treatments. A group of genes were upregulated in both single Sorafenib and combinatory TGX-221 treatments while they are downregulated in single PIK-75 and combinatory PIK-75 treatments. Because of this opposite effect, we anticipated the relation of these genes with antagonistic action of combined therapy of TGX-221 and Sorafenib. In this group, most of the genes were mitochondrial pseudo-genes. It is known that mitochondrial dysfunctions are mostly associated with apoptotic resistance and metabolism of tumor cells and one of HCC hallmarks points out the mitochondrial mutations in cancer development. Furthermore, downregulation of enzymes mediating oxidative phosphorylation for TGX-221 and Sorafenib treatment with respect to PIK-75 treatments confers the previous antagonistic nature in Mahlavu cells [27].

### Network based interpretation with Omics Integrator

A traditional way of RNA-seq analysis is to use only DEG sets for gene enrichment analysis, which generally restricts the capture of complete cellular events. However, application of a conventional method to connect DEGs in a network though their known protein-protein interactions can reveal intersecting/hidden regulation patterns. By using Omics Integrator tool, we adapted Prize Collecting Steiner Tree (PCST) algorithm to associate DEGs by adding intermediate genes (or Steiner nodes) aiming the construction of the most optimal gene-to-gene network. As reference network, we converted protein nodes in STRING human protein interaction network to gene nodes. Using Steiner nodes together with DEG sets introduced more specific gene ontologies. Distribution and relations of PCST enriched gene sets were presented in 5 sets Edwards–Venn diagrams for each treatment (Fig. 5A and 5D). NHBE found to be differentially expressed in PIK-75 (ALPHA) and TGX-221 (BETA), and Sorafenib-PIK-75 (SALPHA). LAMP3, SIRPG and CD83 genes are differentially expressed in TGX-221 (BETA), and Sorafenib-PIK-75 (SALPHA). FOSB is differentially expressed in TGX-221 (BETA), Sorafenib-PIK-75 (SALPHA), and Sorafenib-TGX221 (SBETA). RICTOR and STK32A are differentially expressed in Sorafenib-TGX-221 (SBETA) and Sorafenib (SOR) treatments.

Although, DEG lists found to be highly correlated, network based functional analysis revealed that kinase inhibitor treatments in both cell lines resulted with different biological processes. A better comparison of the networks was provided through a functional encolouring and sizing of the nodes and a systematic usage of the network centrality measures for clustering. PCST predicted optimal gene-to-gene networks were imported into Cytoscape, gene logFC values were used to color the nodes to represent up- and downregulated branches. We arranged the sizes of the nodes according to their betweenness centrality to better organize hub genes. PCST optimal input parameters (final network statistics) were summarized in Supporting results table 3. In Fig. 5B and 5E, network nodes including both DEGs and Steiners were compared. As a result of PCST, hidden expression patterns were identified. Since input DEG numbers for TGX-221 treated Huh7 and Mahlavu cell and Sorafenib treated Mahlavu cell were low, their networks were smaller than the others. Sorafenib and TGX-221 combined treatments in Huh7 and single PI3K-*α* inhibitor (PIK-75) and combined treatments in Mahlavu were separated from the other treatments in the dendrograms.

**Figure 5.**
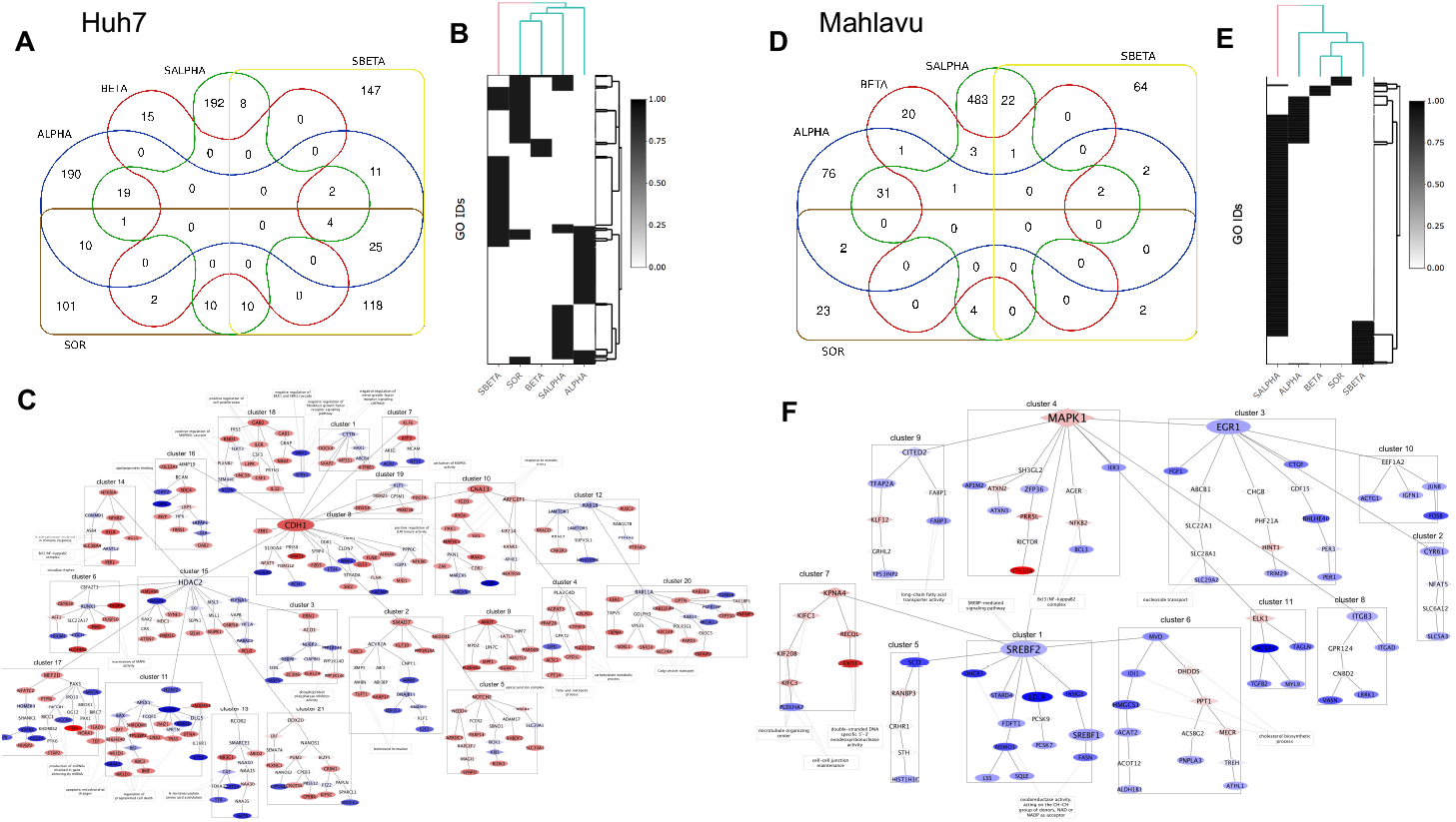
Network based interpretation of DEGs. Venn diagram scheme of Huh7 network nodes **(A)**. Dendogram of GO enrichments for Huh7 **(B)**. Network representation of PI3Ki-*β* and Sorafenib treated Huh7 cells **(C)**. Dendogram of GO enrichments for Mahlavu **(D)**. Venn diagram scheme of Mahlavu network nodes **(E)**. Network representation of PI3Ki-*β* and Sorafenib treated Mahlavu cells **(F)**. ALPHA; PI3Ki-*α* inhibitor, SALPHA; PI3Ki-*α* inhibitor and SOR, SBETA;PI3Ki-*β* inhibitor and SOR, SOR; Sorafenib treatments.

We optimized the networks by limiting the number of trees to one. By minimizing their overall degrees to avoid hairballs we had more than one central hub nodes in the network generating more branches for the analysis (Fig. 5C and 5F). In order to understand the relatedness between the gene nodes in the networks, the functions of the branches should be exploited. Yet, the significance of biological enrichment analysis highly depend on the input size. The power of statistical analysis is low for large DEG sets. Eventually, we clustered networks using gene nodes’ betweenness centralities by Glay algorithm creating branches and applied BINGO for each cluster/branch to get their enriched gene ontologies. Finally, we selected the significant gene ontologies for clusters, and connected them through the network. The ultimate network visualizations allowed us to analyze overall effect of up- or downregulation of genes and provided a comprehensive space for network comparisons through clusters. Other network representations can be reached in the supporting results and cytoscape files are in the referred Cansyl github repository.

### Selection and validation of genes from optimal networks based on centrality metrics

In order to reveal candidate genes to be target for drug studies, optimal PCST networks were analyzed with their centrality metrics. Since hub nodes in the optimal networks were mostly the well-studied genes, we decided to eliminate them to find novel targets in branches. Although, Omics Integrator scales the optimal networks avoiding hub node bias, we also filtered out the nodes that has betweenness centrality values higher than 0.001 after filtration of random nodes (frequency ≥ 0.01).

Then, we used centrality properties of optimal networks which were calculated by Networkx python library. Each network was filtered by degree, eigenvector and betweenness centralities that higher than 0.001 allowing us both not to select the nodes at the end of the branches. The remaining nodes were sorted by inhibitor treatments and eigenvector centrality, and at most six genes were selected for each treatment (Fig. 6A). In the final sets, we have come up with 20 genes for each cell line (Fig. 6B). For Huh7 cells: CDC27, CCDC80, AARS2, ACSBG2 and CITED2 genes in PIK-75 inhibitor treatment, RIMKLA in TGX-221 treatment, CEBPB, DNAJC10, DLK1, EDEM1, ATP6V1D and DUSP8 genes in PIK-75 and Sorafenib combined treatment, LIN7C gene in TGX-221 and Sorafenib combined treatment, EXOC7, FEZ1, GAB2, HOXA10, BIRC7 and ANKRD28 genes in Sorafenib inhibitor treatment and for Mahlavu cells: ATP1B1, CACNA1H, CAPNS1, CCT7, ATG9A and BOLA2B genes in PIK-75 treatments, CGA and TNFRSF4 genes in TGX-221 inhibitor treatment, ALMS1, AOX1, BCL3,CD276, ANKRD1 and ASIC1 genes in PIK-75 and Sorafenib combined treatment, HMGCS1, GDF15, AGER, FABP1, ACOT12, CRHR1 genes in TGX-221 and Sorafenib combined treatment were prioritized for further investigations. For PTEN deficient Mahlavu cells along with single Sorafenib treatment, our prioritization strategy found no significant genes. Here, it is interesting to find some of the targets from Steiner nodes (white boxes) since they cannot be exploited using classical differential expression analysis.

**Figure 6.**
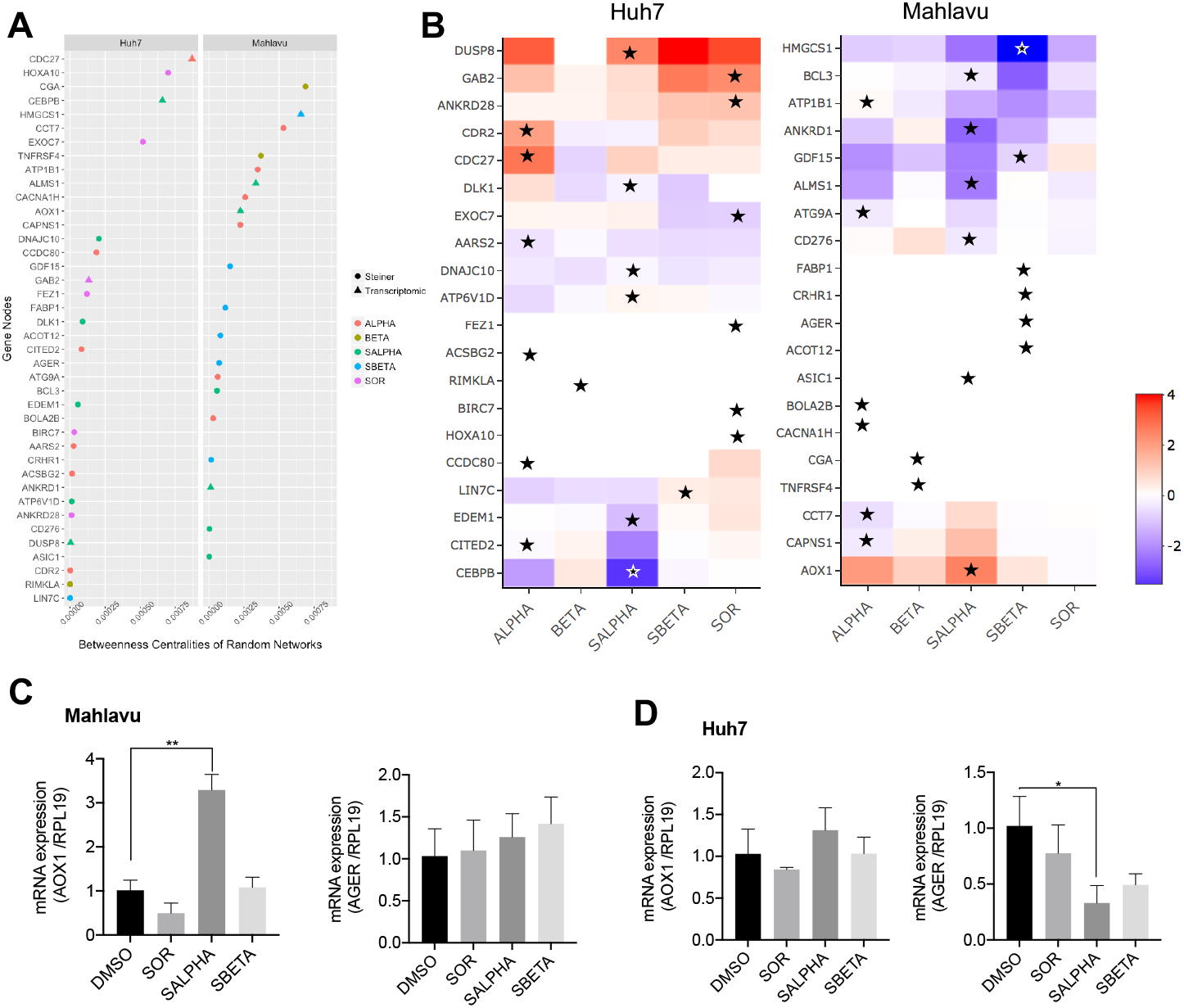
Proposed drug target genes. Prioritized nodes for Huh7 and Mahlavu was ranked by betweenness centrality values of randomized networks for each inhibitor treatment **(A)**. Expressions of the genes in the cell lines. Prioritized treatments were pointed for corresponding drug treatment target **(B)**. Relative expression profile of Mahlavu **(C)** and Huh7 **(D)** cells for AOX1 and AGER genes determined by qRT-PCR. Expression values were normalized with RPL19. Experiment was performed as triplicates and for statistical analysis, unpaired t-test with Welch correction was performed. *p<0.05; **p<0.01. ALPHA; PI3Ki-*α* inhibitor, HBETA; PI3Ki-*β* inhibitor, SALPHA; PI3Ki-*α* inhibitor and SOR, SBETA; PI3Ki-*β* inhibitor and SOR, SOR; Sorafenib treated cells.

We selected AOX1 and AGER genes to be validated by mRNA expression though qPCR experiments (Fig. 6C and D). Mahlavu and Huh7 cells were treated with Sorafenib or its combinations with PI3K-*α* and *β* inhibitors. The gene AGER is selected because it is a pure Steiner node and not found in our DEG list. Whereas AOX1 is in both Steiner node and part of the DEG list. Hence, our qPCR results correlate and validates our network analysis results.

## Discussion

Recently the influence of isoform diversity on responses to drugs with respect to large number of GPCR receptors is demonstrated at systems level [17]. Current studies focusing on the discovery of agents against HCC aim to identify target proteins that escape from regulatory signaling mechanisms of the cell. Conventionally, these studies concentrate on a single gene or locus which result in a comprehensive investigation of a new tumor driver gene, yet, in most cases single driver gene analyses are inadequate to solve the complex network of cancer pathogenesis as the interaction of several signaling pathways and their interlaced connections including HCC which represents high rate of tumor heterogeneity.

PI3K/Akt/mTOR signaling is involved in cellular growth, proliferation and cell cycle progression in various cancer cells [6]. Frequent mutations and loss of function in PTEN tumor suppressor gene leads to constitutive activation of Akt protein hence activation of PI3K/Akt/mTOR cell survival pathway. Sorafenib, which is the most studied therapeutic agent for HCC, targets Raf/MEK/ERK cascade which is compensated by PI3K/Akt signaling activation in favor of cell survival in cancer cells which is one of the reasons for limited effectiveness of this drugs [8]. Sorafenib treatment frequently results in resistance to the treatment within nearly 6 months due to release of pro-inflammatory cytokines and chemokines in tumor microenvironment which promotes cancer stemness, tumor proliferation, and angiogenesis [28, 29]. Hence signaling pathways activated by these molecules in favor of Sorafenib resistance must be targeted in order to alter resistance toward Sorafenib.

There are several drugs in clinical trials targeting PI3Ks inhibiting tumor progression [22]. PI3K inhibitors are classified as pan-PI3K inhibitors and isoform-specific inhibitors. Current molecular and clinical trial studies focus on the effectiveness of these inhibitors as well as the mechanisms of resistance to PI3K inhibition. In this study, we investigated the molecular alterations in combinatory treatment of isoform PI3kinase inhibitors targeting PI3K/Akt/mTOR pathway with Raf/MEK/ERK signaling inhibitor Sorafenib either on PTEN adequate (Huh7) or deficient (Mahlavu) HCC cells (Fig. 1). PI3K inhibitors usually combined with mTOR inhibitors in order to increase the effectiveness of the treatment [22, 30]. However, there are limited studies which targets alternate cell signaling survival pathways (i.e. Raf/MEK/ERK vs PI3K/Akt/mTOR) with the aim of reveling the genes involved in synergistic or antagonistic resistance to inhibitors.

We showed a synergistic inhibition of cell growth in both cell lines treated with PI3K-*α* inhibitor PIK-75 and Sorafenib. The synergistic cytotoxicity was more effective in PTEN-adequate Huh7 cells. While combinatory PI3K-*β* inhibitor TGX-221 treatment also synergistically inhibit cellular growth in Huh7, we observed a strong antagonistic effect in Mahlavu cells indicating the importance of isoform specific actions of Phosphatidylinositol 4,5-bisphosphate 3-kinase catalytic subunit isoform kinase inhibitors. We then investigated the molecular mechanisms involved in this differential phenotypic response to isoform specific inhibition by comprehensive network analysis of RNA-seq data upon treatment with drugs in combination with Sorafenib. Both inhibitors resulted with up regulation of key pathways in inflammation and immune response like; BCL3/NF-*κ*B, cell proliferation (CDH2 and CCD1), Jun kinase and osmotic stress in Huh7 cells. Moreover, while in TGX-221 and Sorafenib treatment, genes having role on regulation of programmed cell death and apoptotic mitochondrial changes were downregulated. It is known that interactions of GTPases with PI3K are isoform specific, while Ras cannot bind to p110-*β*, RAC1 and CDC42 proteins can activate p110-*β* [31]. Our network analysis showed that PI3K-*α* inhibitor (PIK-75) and its combined treatment with Sorafenib also showed a correlation in Huh7 cells. They both resulted with negative regulation of Erk1/Erk2 signaling and activation of MAPKK activity. Combinatory treatment with Sorafenib, mostly resulted with downregulation of hub proteins JUN, INSIG1, MDM2 and SOX9 associated with cancer cells. Hence, in PTEN adequate Huh7 cells, PIK-75 and Sorafenib treatment could decrease cell proliferation and decrease Sorafenib dependent immune response. Our data indicates that targeting PI3Ki-*α* isoform in an inhibited MAPK pathway background with Sorafenib would be a better therapeutic approach in both PTEN deficient and adequate Hepatocellular cancer cells.

However, combination of PI3K-*β* inhibitor (TGX-221) and Sorafenib in Mahlavu cells showed a strong antagonistic action, which probably depends on PI3Ki-*α* isoform activity. Our data with PTEN deficient Mahlavu cells demonstrated that constitutive PI3K/Akt pathway activation makes these cells more resistant due to on PI3K-*α* isoform activation since the inhibition of PI3K-*β* with its specific inhibitor TGX-221 makes these cells to resistant to Sorafenib. When TGX-221 and Sorafenib was combined, MAPK and nuclear factor kappa B (NF-*κ*B) signaling upregulated and increasing activity in response to immune stress and inflammatory injuries genes were enriched (Fig. 4 and 5). In Mahlavu, TGX-221 combined with Sorafenib treatment shows a decreased level of Bcl-3 responsible of antagonistic action.

In this study we also perform system level network analysis in order to identify genes involved in isoform specific actions of PI3K inhibitors using DEG genes from our RNA-seq transcriptome data on STRING human protein interaction networks. Our prioritization strategy using topological features of the optimized networks identified hub and Steiner nodes representing genes involved in differential synergistic or antagonistic effects of isoform specific PI3K-*α* (PIK-75) or PI3K-*β* (TGX-221) inhibitors with Sorafenib. Many of these (DLK1, GAB2, BOLA2B, AOX1 and AGER) closely related to cell proliferation and tumor progression stimulates cell proliferation and associated to poor prognosis in HCC [32, 33, 34]. Steiner nodes prioritized in Mahlavu cells treated with PI3K-*α* (PIK-75) and Sorafenib identified Aldehyde oxidase 1 (AOX1), and Mahlavu cells treated with PI3K-*β* (TGX-221) and Sorafenib identified Advanced glycosylation end product-specific receptor (AGER) genes. Differential expression of these genes are validated by qPCR experiments as shown in Fig. 6C and D. Both genes are associated with glucose metabolism and generation of reactive oxygen species and are involved in proinflammatory actions in liver carcinogenesis

## Conclusion

Combination of targeted drugs to inhibit alternative compensatory pathways holds great promise for effective treatment of cancer including HCC. As we clearly demonstrated in this study system level analysis of cellular networks in response to combination treatments and the investigation of the regulation signaling pathways are of necessity, because such treatments may result in an opposite of the desired effect. The importance of context dependent (PTEN status) PI3K/Akt/mTOR signaling inhibition must be taken into consideration during the use of isoform specific or pan-PI3K inhibitors in combination therapies with Sorafenib with respect to resistance in HCC cells.

## Material and Methods

### Cell lines and kinase inhibitors

Mahlavu and Huh7, HCC cell lines were cultured in DMEM medium, supplemented with 10% fetal bovine serum (FBS), 1% penicillin/streptomycin (P/S) and 1% non-essential amino acids (NEA) and incubated in humidified 37°C incubator with 5% CO2.bMahlavu and Huh7 cell lines were treated with the inhibitors which are listed in Supporting result table 1 (Sorafenib (Nexavar) was from Bayer Health-care Pharmaceuticals, Inc., NJ USA, Inhibitors PIK-75 (cat#528116), TGX-221 (cat#528113), LY294002 (cat#440202) were from Calbiochem).

### Cytotoxicity and cell cycle

Huh7 (2000cell/well) and Mahlavu (1000cell/well) cell lines were seeded into 96-well plates in 150*μ*l of medium/well. The next day, cells were treated with the drugs (Sorafenib (Nexavar) Bayer Bayer Healthcare Pharmaceuticals, Inc., NJ USA, or PIK-75 (PI3K-*α* inh.), TGX-221 (PI3K-*β* inh.), LY294002 (PanPI3Ki) and control (DMSO) in triplicates. After 72h, the media was discarded, and the wells were washed with PBS, and 50*μ*l of 10% cold TCA (Merck, Germany) was added for fixation and incubated with TCA at +4°C in dark for 1 hour. Then cells were washed with ddH_2_O for 4 times, and the plates were air-dried at room temperature. Finally, 50*μ*l of 0.4% sulphorhodamine B (SRB) (Sigma Aldrich) solution in 1% acetic acid was applied to each well, and the plates were incubated for 10min in dark at room temperature. Excess dye was washed off with 1% acetic acid (4 or 5 washes). Finally, 200*μ*l of 10mM cold Tris-Base was applied to each well to solubilize SRB. Then, the absorbance values were measured at 515nm and were analyzed to determine the effect of each drug on cell proliferation compared to control [37]. Sorafenib, PIK-75 and TGX-221 were used to treat HCC cells in concentrations which described in respective figure legend and the cells were incubated for 96 hours. Cell viability and DNA content calculations in flow cytometric cell cycle analysis was performed using propidium iodide.

#### Real-time cell electronic sensing (RT-CES) system for cell growth and cytotoxicity analysis and synergy analysis

50*μ*l Huh7 (2000 cell/well) and Mahlavu (1000 cell/well) cells were seeded into 96-well plates in 100*μ*l of medium/well. The next day, cells were treated with the drugs and CI (Cell Index)values were taken every 10 min for 4h to get the fast drug response and then every 30 min to obtain the long-term drug response. Impedance measurements displayed as CI values reflect cell growth. The SynergyFinder Zero Interaction Potency (ZIP) model is used for the evaluation of the combined effect of PIK-75, TGX-221, and their combinations with Sorafenib (Sorafenib (Nexavar) Bayer Healthcare Pharmaceuticals, Inc., NJ USA). ZIP model defines the effect of combining two compounds by comparing the change in the dose-response curves between individual drugs and their combinations [38].

#### RNA extraction and sequencing

Total RNA was isolated with NucleoSpin RNA II Kit (Macherey-Nagel) according to the manufacturer’s protocol (MN, Duren, Germany) with small modifications such as 30 min of DNA digestion instead of 15min and 2-step elution with 20*μ*l water instead of one elution with 60*μ*l. RNA concentration was measured with NanoDrop and A260/A280, A260/A230 ratios were checked for RNA quality and purity. Total RNAs were provided to BGI Tech (https://en.genomics.cn/) for sequencing. RIN values are acquired Agilent Bioanalyzer system and they were above 0.8 for all samples. Details of RNA-seq experiment and data can be found at PRJNA556552.

#### Wound Healing

In the wound-healing assay, a wound was made in the middle of a confluent cell monolayer and the migration of cells to this area was assessed by taking photos at different time points and calculating the wound closure with respect to the initial wound width. Sorafenib, and TGX-221 were used at a concentration of 10*μ*M, except PIK-75 inhibitor, which was used at 0.1*μ*M concentration. Photos of the wounds were taken after 24h and 48h. The sizes of the wounds were calculated at all time points. At least 12 different wound distances were noted for each condition at each time point and the averages were used for analysis to construct the graphs.

#### Quantitative RT-PCR (qRT-PCR)

Mahlavu (100.000 cells/dish) and Huh7 (250.000 cells/dish) were seeded into 10 cm culture dishes. After 24 hours, cells were treated with the inhibitors and incubated for 48 h and then collected for RNA isolation. RNA-purification kit (Qiagen, cat#74106) and cDNA synthesis (ThermoFischer, cat#K1621) kit were used according to manufacturer’s protocol. Total RNA amounts were measured with Nanodrop One (ThermoFisher). qPCR was initiated with 50ng cDNA and performed with FastStart Essential DNA Green Master (Roche, cat#6402712001) via Roche LightCycler 96 Instrument, according to manufacturer’s protocol optimized for this instrument. Primer sequences are: AOX1-f: 5’-ggggtgttccgtgtttttcg-3’, AOX1-r: 5’-caggttcatctctcggaatcattt-3’, AGER-f: 5’-agcatcagcatcatcgaacca-3’, AGER-r: 5’-gc-ctttgccacaagatgacc-3’ and RPL19-f: 5’-gctctttcctttcgctgctg-3’, RPL19-r: 5’-ggatct-gctgacgggagttg-3’. All reactions were performed in triplicates. The Ct (cycle threshold) values were normalized against RPL19 reference gene

### Bioinformatics Methods

#### RNA-Seq analysis

RNA reads were processed by Illumina Hiseq 2000 (SE50). 12 FASTQ files (PRJNA556552), were first analyzed through a well-known quality assessment tool; FASTQC [40]. Then, without any trimming, single-end reads were aligned to the reference human genome (GRCh38/hg38) using a split read aligner algorithm TopHat V2.1.0 [41]. TopHat itself features an ultrafast mapper Bowtie v2.2.6 algorithm [42]. After that, aligned reads were quantified by HTSeq-count v0.6.1 [43] for given human gene split regions (GRCh38 v84) to count how many transcripts map to each gene, which generates a gene level count matrix.

FASTQC analysis of this study results are included into the referred CanSyl github repository and other alignment metrics summarized in supporting results table 2.

#### Differential expression analysis for sequence count data

EdgeR [26], from Bioconductor package, is a widely used method for differential expression analysis. We used gene level count matrices of 12 RNA-seq treatment sets as input of EdgeR. DMSO treated Huh7 and Mahlavu cells were used as negative controls. EdgeR constructs a negative binomial model using the RNA count data. In our experimental design, there was no biological replicates of the samples to inherit the in-sample variation. EdgeR solves no-replication problem by suggesting a different dispersion calculation method to estimate variation within each sample compared to housekeeping genes. A set of housekeeping genes in Hepatocellular carcinoma was well characterized in Ersahin T. et al [44]. We used these housekeeping genes to estimate biological coefficient of variation (BCV) value manually.

Before EdgeR analysis, genes with less than 5 readings were filtered out using counts per million constraint (cpm ≤ 5). A biological model was constructed by taking BCV as 0.045. Differential analysis performed using *exactTest* function of EdgeR package. Then, we limited logFC (log2 of fold change) to −/+2 for both cell lines. However, with this limitation, Mahlavu had no significant numbers of DEGs for downstream analysis. On the other hand, Huh7 resulted greater number discounting for single PI3Ki-*β* treatment. Therefore, for Mahlavu cell lines, a less stringent logFC value (−1.5/+1.5) was used for further analysis. Finally, we selected the top DEGs according to following filters; p-value ≤ 0.01, FDR ≤ 0.01, and logFC ranges (−2/+2) for Huh7 cell and (−1.5/+1.5) for Mahlavu cell. Gene annotations were obtained using *org.Hs.eg.db R* package [45] from Bioconductor.

#### Dendogram analysis

Heatmap representation is one of the most popular graphical methods for visualization of big-data providing color encoding cells that represent numbers. Heatmaply [46] is a very powerful way of investigating clusters in a high dimensional data since final heatmap result is visualized as interactive graph offering inspection over the cells making possible for zoom-in. In our study, all dendograms were visualized through heatmaply using default *hclust* clustering by using Euclidian as distance measure.

#### Gene Ontology (GO) analysis

Given a set of genes on the network, Cytoscape plug-in BiNGO tool [47] maps functional terms to enriched genes to output GO terms and their statistical features. In order to have a better understanding of the processes that selected genes having role on, statistically over-represented GO terms were characterized using BiNGO in our analysis. We have used a very stringent Benjamini&Hochberg False Discovery Rate (FDR ≤ 0.005) to filter out non-significant GO-terms. GO sets containing redundant and electronically annotated terms generated noise for functional comparisons. In order to avoid those suspicious GO terms, we have only used GOs with experimentally validated codes (EXP, IDA, IEP, IGI, IMP, and IPI). The codes were matched though Gene Ontology Annotation (GOA) database.

#### Network construction and optimization of DEGs using PCST approach

PCST (Prize Collecting Steiner Tree) [48, 49] aims to identify sub-networks from an interaction network given a set of weighted genes. By using PCST, we have extracted the biologically meaningful interactions between the DEGs from human protein-protein interaction data. We used Omics Integrator software to implement PCST algorithm. We used Forest module in our analysis to determine multiple sub-pathways in the human interactome. PCST algorithm finds an optimal tree, including the terminal nodes (from DEG lists in our case) with prizes travelling through the interactome nodes which have costs of edges only if they are included. The task is to find the shortest paths between the prize nodes avoiding the costs on the edges. The algorithm minimizes the cost of all edges by passing through as much prize nodes possible. In order to construct meaningful trees using DEGs, forest parameters must be fine-tuned. The size and degree of the forests are expected to vary as the number of genes in the input files changes. Forest parameters depend highly on the distribution of prizes and numbers of the nodes. The best combinations of parameters for each DEG set was explored using forest-tuner [50] which is PCST algorithm parameter tuner for *ω*, *β* and *μ* parameters. This script was used to find the best arrangements of the parameters to be used in Forest module for each treatment. We had searched the parameters in the following ranges: *ω* (1-10.0 or 5-15), *β* (1-15.0), *μ* (0.01-0.05). Here, *ω* parameter tunes the number of trees in the network, *β* parameter increases the number of prices entering the tree and *μ* is another parameter that arranges the dominance of hub proteins in the network. Among all of the possible solutions, we have selected the combination which generates a network with minimum mean degree. Optimal PCST parameters are summarized in supporting results table 3.

The interactome set given to Forest module was derived from STRING protein-protein interaction database v10 [51]. In STRING, network edges were scored according to a confidence score (range of 0 to 1) determined through an algorithm by the database. The confidence score gets higher as it gets more experimental proofs basically. In our analysis, we used the interactions only with high confidence proofs (at least 0.7). Then, we applied *−log10* conversion to the confidence score, to use them as edge costs in Omics Integrator.

#### Randomization tests

In order to test the significance of the nodes appearing in the optimal nodes, each PCST network was subjected randomization tests using forest module (*–randomTerminals 100*). The tests were performed using random set of terminals with respect to keeping node numbers, and original interactome set with same edge weights and optimization penalties the same. The probability that a node randomly to be connected in the network was expressed by its frequency of randomness in the network. Therefore, less frequent nodes would be the most specific ones to the network. Through the analysis, we had used nodes appeared only once in the random networks.

#### Network centrality

Centrality measures are the indicators of most valuable vertices in the graph for network analysis and they are often used to identify influential nodes of the network providing a ranking which identifies the important nodes in the network. We had used degree, eigenvector and betweenness centralities in order to estimate network topology. Networkx python library [52] was used to calculate centrality measures.

#### Effective visualization and clustering of the networks

Omics Integrator Forest module generates networks in *.sif* format which is compatible format for Cytoscape visualizations. Cytoscape includes many add-on for biological network analysis, therefore we both analyzed and visualized our networks on this tool. For our study, after *.sif* file was imported into the Cytoscape yFiles layout algorithms was implemented and hierarchical layout was selected for visualizations. In order to cluster the networks, Glay algorithm [53] using edge betweenness centrality is implemented. For proper annotation of the clusters AutoAnnotate plug-in [54] was used. Application of this strategy resulted with the most connected patterns in the networks. Then, in order to better understand what processes of these patterns having role on, statistically over-represented Gene Ontology (GO) terms were characterized using BiNGO for each cluster in networks. Selected GO terms were also imported into the network using their gene maps. The final visualizations represented all gene relationships, up- and downregulated genes and internal Steiner nodes. Also, highly connected groups and GO annotations were provided for an easy and efficient way to compare networks with each other.

#### Prioritization of nodes in PCST generated networks

Based on the network topology, we developed a prioritization strategy for further investigation as drug targets. A node is treatment specific only if it occurs in the branches on random networks while present in more central areas on the optimal networks. In order to accomplish these nodes, we used the least frequent nodes (0.01) resulted from randomization test. Here, hub nodes of optimal networks have selected though using degree, eigenvector and betweenness centralities greater than From these nodes, we eliminated the ones that were predominant in the random network using degree centrality of random networks smaller than 0.001. Finally, top 20 nodes for each treatment were selected and represented.

## Supporting information

supporting results

## Abbreviations

HCC: Hepatocellular carcinoma
DEGs: Differentially expressed genes
GO: Gene Ontology
FDR: False Discovery Rate
logFC: log2 of fold change

## Availlabiliy of data and materials

FASTQ reads of the RNA-seq experiments are available in the NCBI-SRA repository [https://www.ncbi.nlm.nih.gov/sra/PRJNA556552]. The datasets generated and analysed during the current study are available in the CanSyL lab github repository [https://github.com/cansyl/Isoform-spesific-PI3K-inhibitor-analysis].

## Acknowledgements

We are grateful to our colleagues part of CanSyl laboratory for their commends on the study and manuscript.

## Contributions

KN analyzed the data and write the manuscript. DCK and TE performed the experimental analysis. DCK write the manuscript. AK performed the qRT-PCR experiments. TE performed the RNA-seq experiments. NT provided Omics Integrator tool and helped through data analysis. RCA designed the experiments and write the manuscript.

## Ethics declarations

### Ethics approval and consent to participate

Not applicable

## Consent for publication

Not applicable

## Compering interest

The authors declare that they have no competing interests.

## Additional Files

### Supporting results

IC50 values, RNA-seq quality,optimal PCST networks and clustering and enrichment analysis of PCST generated networks listed and figured in this additional file.

