## supporting results for "Context dependent isoform specific PI3K inhibition confers drug resistance in Hepatocellular carcinoma cells"

### Supplementary Results

#### $IC_{50}$ values

$IC_{50}$ values at 72h calculated for both Sulforhodamine B (SRB) cytotoxicity assay and Real-time cell electronic sensing (RT-CES) system shown in Table 1.

#### RNA-seq quality

12 Single-end RNA reads, processed by Illimuna Genome Analyzer, were analyzed using FASTQC. According to FASTQC reports, length of 49bp single-end reads were well sequenced and no end bias was seen for the reads. An average of 33,465,904 single end 49bp clean reads was generated. The average mapping rate was 97.28%, resulting an average coverage of depth 41X (Table 2). Then, the mapped reads were counted using HTSeq-count given a set of genomic features. Library sizes were calculated through the gene set. Without any transformation on to counts, library sizes of genes were between 28-38 thousand.

#### Optimal PCST Networks

Forest-tuner was run for DEG results to find the best arrangements of parameters in this ranges; $\omega$ (1-10.0 or 5-15), $\beta$ (1-15.0), $\mu$ (0.01-0.05). From the possible solutions, we selected the networks with the smallest mean degrees which was listed in Table 3. PCST networks were analyzed further through heatmap and cytoscape visualizations.

#### Clustering and enrichment analysis of PCST generated networks

Cytoscape visualizations of inhibitor treated Huh7 and Mahlavu networks were presented in Figure 1 to 10. The networks were generated through PCST algorithm using Omics Integrator and sketched by Cytoscape tool. The nodes were colored by logFC values resulted from edgeR analysis and the darkness of the color increases by the increase of the value. Red and blue represented up- and downregulation respectively. Steiner nodes were shaped as diamond while input transcriptome nodes were figured as ellipse. Node size was directly correlated with betweenness centrality of nodes. Clusters were generated using betweenness centralities of the nodes using a community cluster algorithm (Glay). Clusters were boxed for a better representation. Then, the clusters were separately analyzed by BiNGO for gene set enrichment analysis and only selected significant Gene Ontology was added to the network tough associated genes.

### Tables

**Table 1 $IC_{50}$ values at 72 hours of incubation, calculated based on SRB and RT-CES assays.**

| | SRB | | RT-CES | |
|---|---|---|---|---|
| Inhibitors | Huh7 | Mahlavu | Huh7 | Mahlavu |
| Sorafenib | 8.0$\mu$M | 6.6$\mu$M | 10.0$\mu$M | 10.0$\mu$M |
| PIK-75 | $\leq$0.3$\mu$M | $\leq$0.3$\mu$M | 0.1$\mu$M | 0.1$\mu$M |
| TGX-221 | $\geq$40.0$\mu$M | $\geq$40.0$\mu$M | 10.0 $\mu$M | 15.0$\mu$M |
| LY294002 | 3.8$\mu$M | 8.7$\mu$M | 10.0 $\mu$M | 10.0$\mu$M |

**Table 2 Total sequences processed, map rate average sequence length and GC% content and coverage of the experiment.**

| | Total sequences | Map Rate(%) | Sequence Length | GC Content(%) | Coverage* |
|---|---|---|---|---|---|
| Akt Normal (Huh7) | | | | | |
| PIK-75 | 33192992 | 97.0 | 49 | 48 | 41 |
| TGX-221 | 32730853 | 97.3 | 49 | 50 | 40 |
| PIK-75 + Sorafenib | 34017394 | 97.4 | 49 | 49 | 42 |
| TGX-221 + Sorafenib | 28579376 | 97.6 | 49 | 49 | 35 |
| Sorafenib | 36506606 | 97.7 | 49 | 49 | 45 |
| DMSO | 37827459 | 97.5 | 49 | 48 | 47 |
| Akt Hyperactive (Mahlavu) | | | | | |
| PIK-75 | 34302640 | 96.0 | 49 | 47 | 42 |
| TGX-221 | 29776671 | 97.5 | 49 | 49 | 37 |
| PIK-75 + Sorafenib | 31897951 | 96.8 | 49 | 49 | 39 |
| TGX-221 + Sorafenib | 37221329 | 97.7 | 49 | 49 | 46 |
| Sorafenib | 32116151 | 97.4 | 49 | 50 | 39 |
| DMSO | 33421422 | 97.5 | 49 | 49 | 41 |

$*Coverage = SequenceLengthSequencesTranscriptomeLength(39841315)$

**Table 3 Selected parameters for PCST analysis using forest-tuner and numbers of nodes, terminals and prizes of generated networks.**

| | $\omega$ | $\beta$ | $\mu$ | (Terminal+ Steiner) Total Node | Prize Node | Mean degrees |
|---|---|---|---|---|---|---|
| Akt Normal (Huh7) | | | | | | |
| PIK-75 | 7.75 | 5.50 | 0.04 | (138+124) 262 | 171 | 24.83 |
| TGX-221 | 5.50 | 3.25 | 0.03 | (5+12) 17 | 5 | 23.08 |
| PIK-75 + Sorafenib | 10.0 | 10.0 | 0.02 | (145+101) 246 | 178 | 28.58 |
| TGX-221 + Sorafenib | 10.0 | 3.25 | 0.03 | (178+147) 325 | 213 | 23.86 |
| Sorafenib | 10.0 | 7.75 | 0.05 | (157+124) 281 | 187 | 27.19 |
| Akt Hyperactive (Mahlavu) | | | | | | |
| PIK-75 | 10.0 | 7.75 | 0.03 | (52+63) 115 | 84 | 29.14 |
| TGX-221 | 7.75 | 5.50 | 0.03 | (6+20)26 | 6 | 16.7 |
| PIK-75 + Sorafenib | 10.0 | 3.25 | 0.01 | (321+236) 547 | 409 | 30.31 |
| TGX-221 + Sorafenib | 10.0 | 5.50 | 0.03 | (53+40) 93 | 75 | 34.65 |
| Sorafenib | 5.0 | 7.00 | 0.04 | (16+15) 31 | 27 | 19.67 |

### Figures

Networks representing differential expression patterns in PI3K/AKT/mTOR and RAF/MEK/ERK signaling pathways in Huh7 and Mahlavu cell lines for differential inhibitory treatments are shown

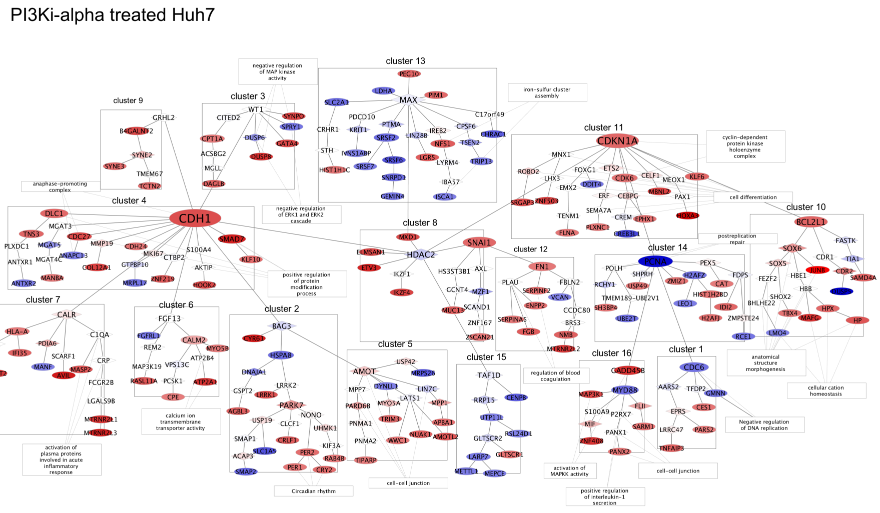


**Figure 1 PIK-75 treated Huh7 cell line network.**

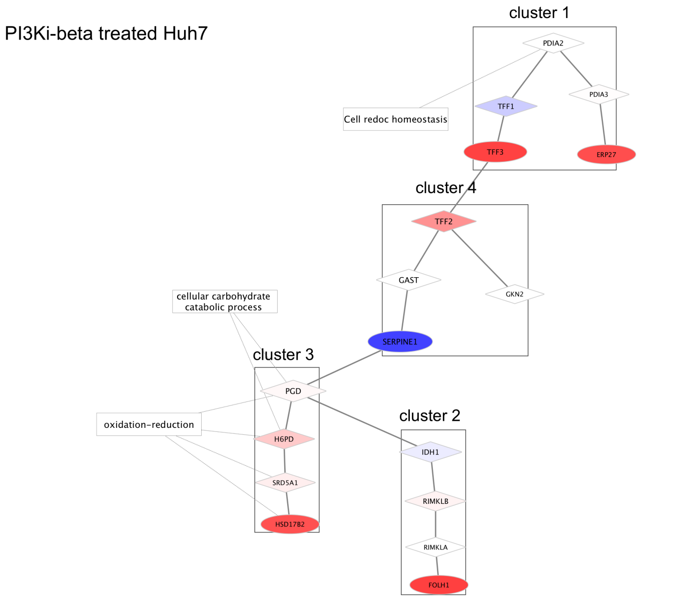


**Figure 2 TGX-221 inhibitor treated Huh7 cell line network.**

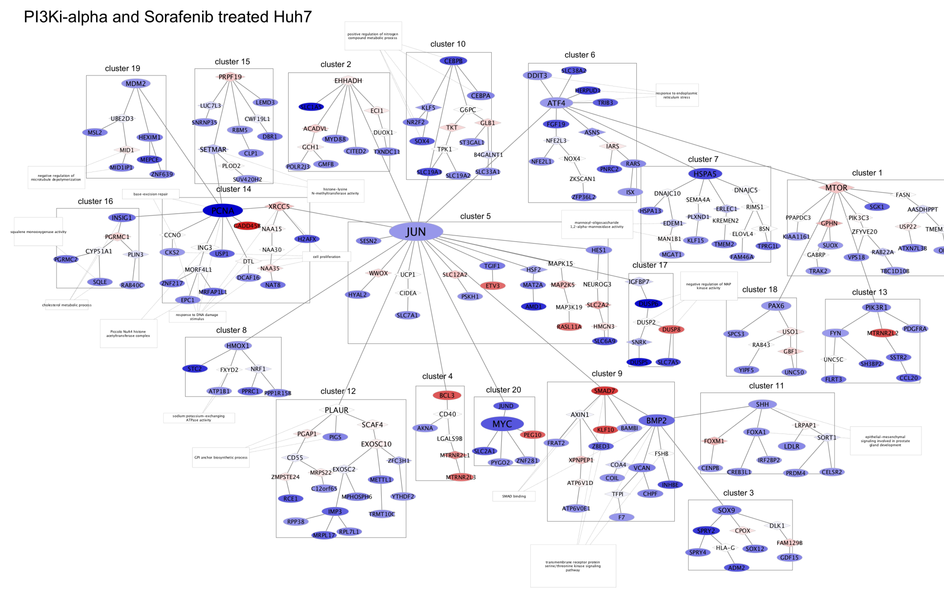


**Figure 3 PIK-75 + Sorafenib treated Huh7 cell line network.**

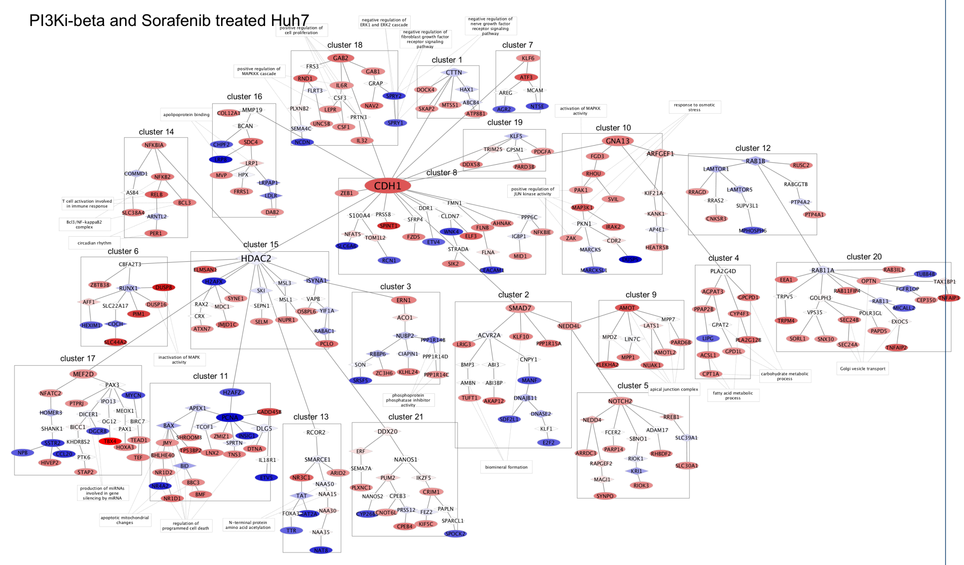


**Figure 4 TGX-221 + Sorafenib treated Huh7 cell line network.**

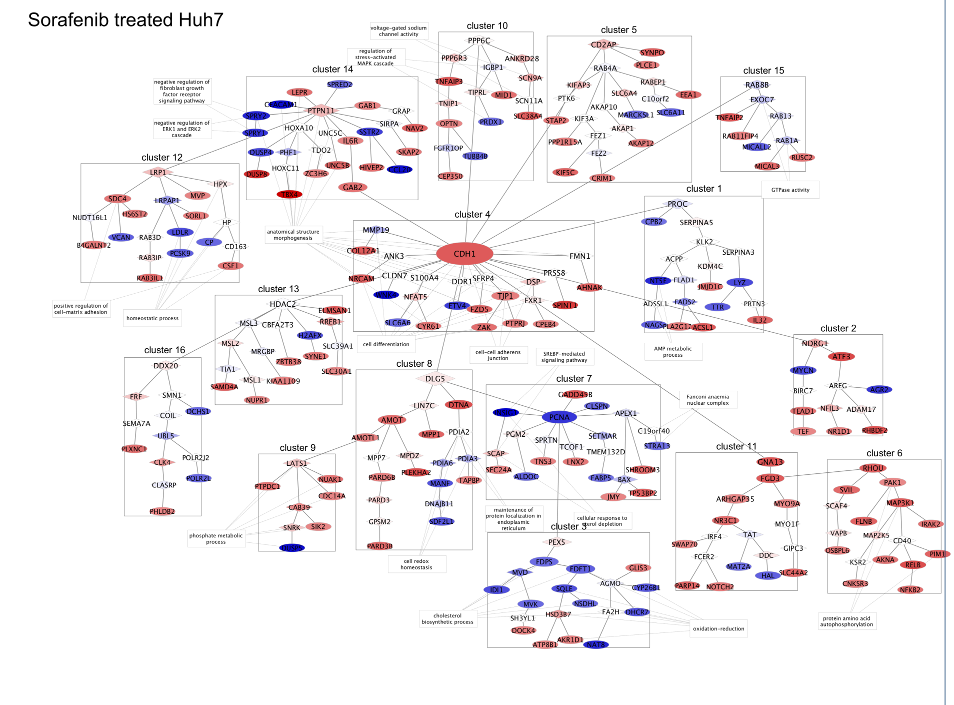


**Figure 5 Sorafenib treated Huh7 cell line network.**

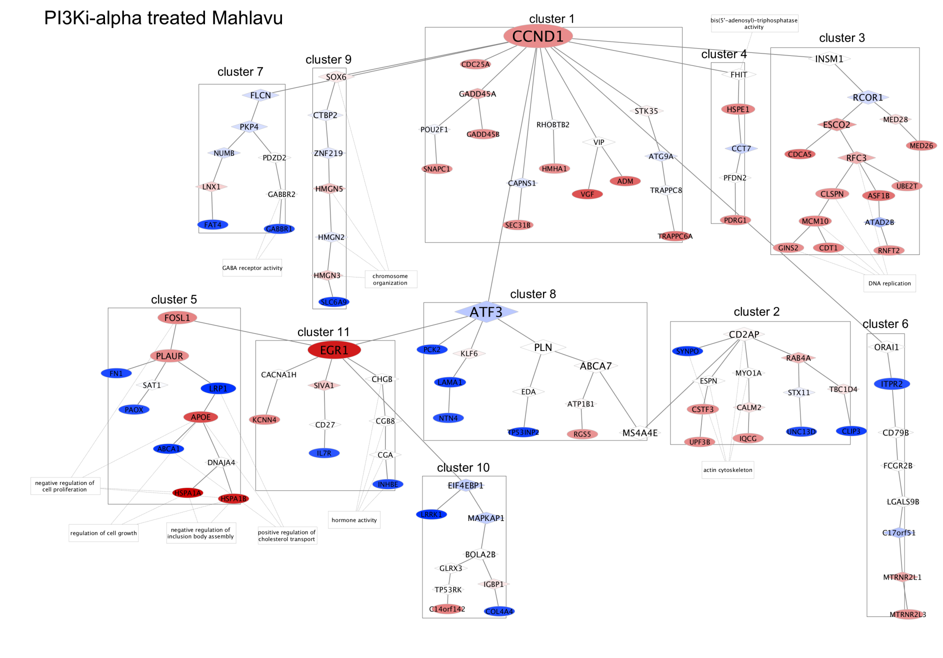


**Figure 6 PIK-75 inhibitor treated Mahlavu cell line network.**

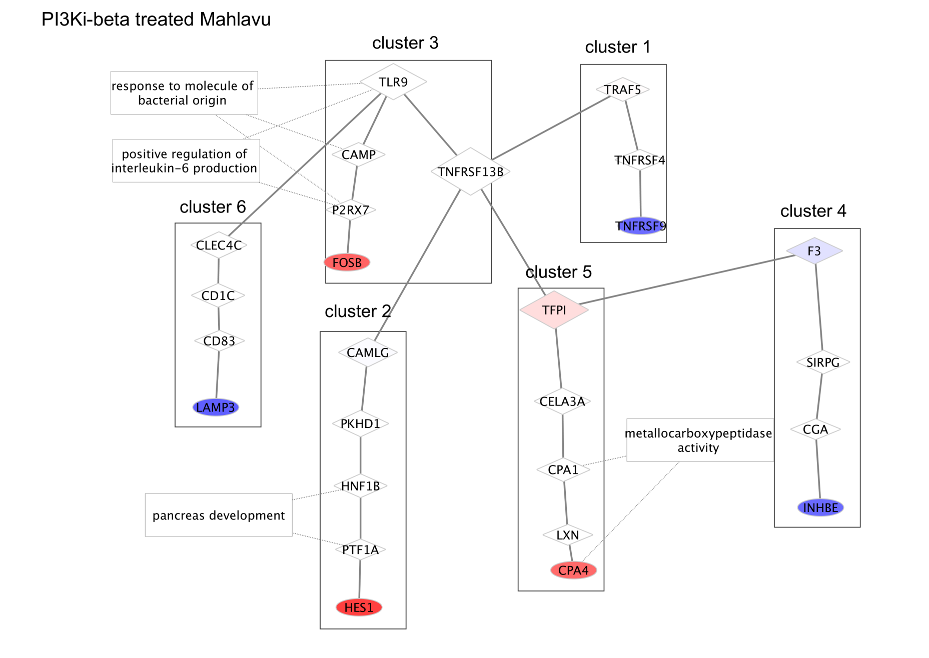


**Figure 7 TGX-221 treated Mahlavu cell line network.**

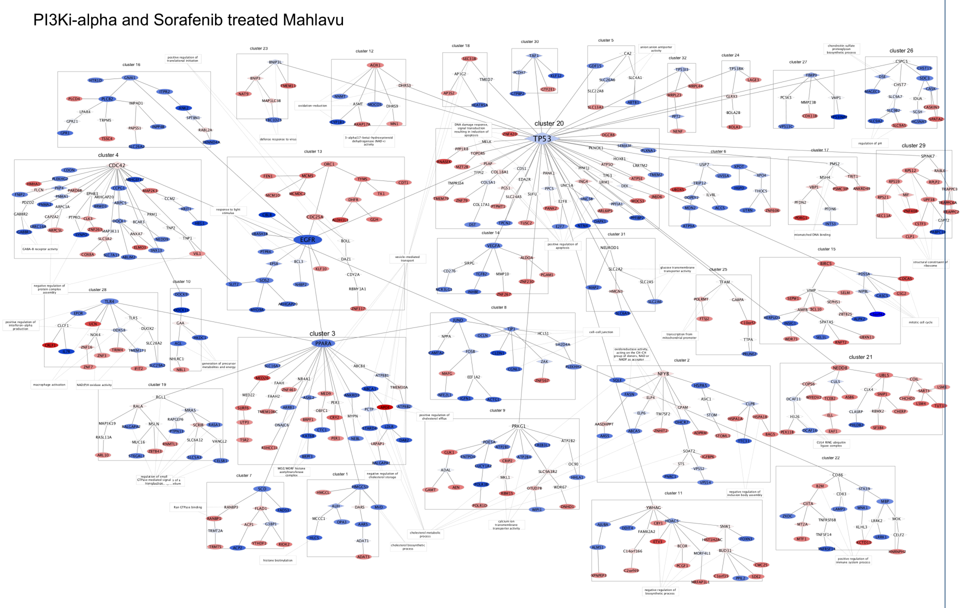


**Figure 8 PIK-75 + Sorafenib treated Mahlavu cell line network.**

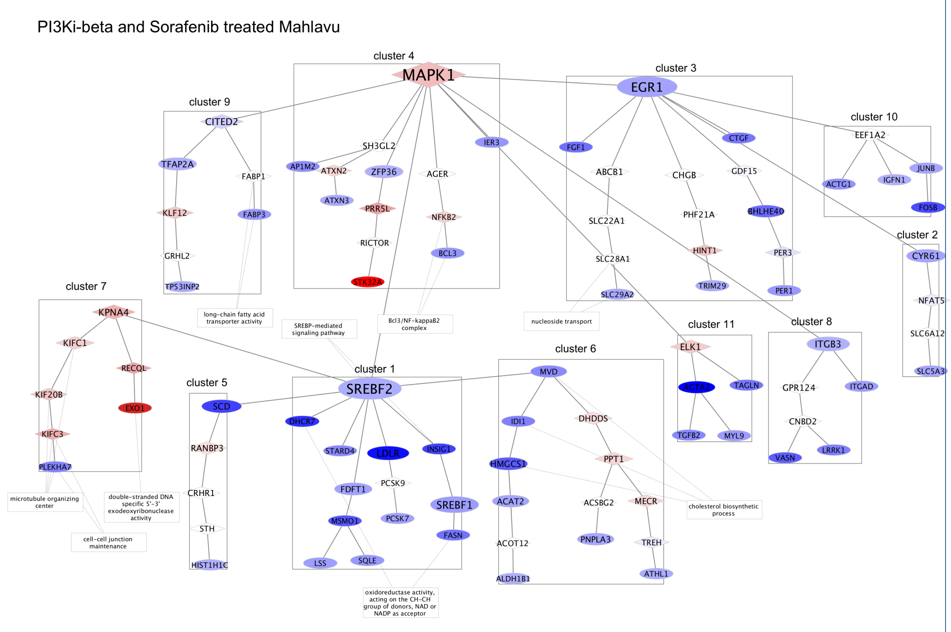


**Figure 9  TGX-221 + Sorafenib treated Mahlavu cell line network.**

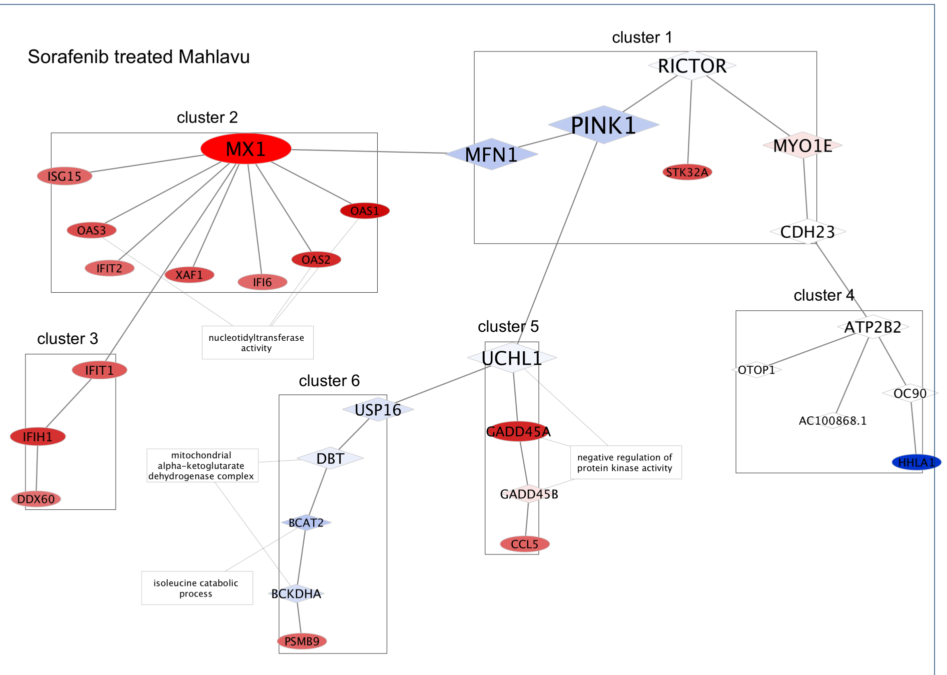


**Figure 10  Sorafenib treated Mahlavu cell line network.**
